# Relationship between the colors of the rivers in the Amazon and the incidence of malaria

**DOI:** 10.1101/2023.06.29.547004

**Authors:** Fernanda Fonseca, Jean-Michel Martinez, Antônio Balieiro, Jesem Orellana, Naziano Filizola

**Author notes:** **Corresponding author:** *Fernanda Fonseca*. **Author Contributions:** Fernanda Fonseca (Conceptualization, Data Curation, Formal Analysis, Funding Acquisition, Investigation, Methodology, Validation, Visualization, Writing – Original Draft Preparation, Writing – Review & Editing), Jean-Michel Martinez (Formal Analysis, Methodology, Supervision, Validation, Writing – Review & Editing), Antônio Balieiro (Formal Analysis, Methodology, Validation, Writing – Review & Editing), Jesem Orellana (Formal Analysis, Methodology, Validation, Writing – Review & Editing), Naziano Filizola(Conceptualization, Formal Analysis, Funding Acquisition, Investigation, Methodology, Project Administration, Supervision, Validation, Visualization, Writing – Review & Editing).

## Abstract

**Background:** Malaria is an infection caused by parasites of the genus Plasmodium, which are transmitted to humans via the bite of the female Anopheles mosquito. In Brazil, approximately 99% of malaria cases are concentrated in the Amazon region. Os rios desempenham um papel importante no ciclo de vida da malária, uma vez que o vetor se reproduz em ambiente aquático. The waters of the rivers in the Amazon have distinct chemical characteristics and this prompted us to analyze the influence of the color of the waters of the rivers in the Amazon on the distribution of malaria.

**Methodology:** This study was conducted for a period of seventeen years (2003-2019) in 50 municipalities in the state of Amazonas, Brazil. A generalized linear mixed model was developed to analyze the association of malaria incidence and three types of river color: white, black and mixed.

**Principal Findings:** The results suggest that there is a trend towards a decrease in malaria cases until 2015, with a possible resumption of the incidence of the disease from 2017 onwards, in all types of river color. In addition, the research indicates that places located near black-or mixed-water rivers have a higher incidence of malaria when compared to places on the banks of white-water rivers.

**Conclusions:** Historically, the hydrological regime has played an important role in the dynamics of malaria in the Amazon, but little is known about the relationship between river colors and the incidence of the disease. In this sense, our results, by showing a significant association between the colors of the rivers and the incidence of malaria over time, seem to expand the understanding between physical-chemical characteristics of the rivers and the occurrence of malaria.

**Author summary:** Malaria is a disease that has a major impact on morbidity in tropical and subtropical countries. In Brazil, most of the cases registered in the country are concentrated in the northern region. A variety of factors can contribute to the occurrence of malaria outbreaks or epidemics, and rivers play an important role in the transmission of the disease. The waters of the rivers in the Amazon have distinct chemical characteristics and this study sought to analyze the influence of the color of the waters of the rivers in the Amazon on the distribution of malaria. Data on the incidence of malaria, from 2003 to 2019, in 50 municipalities in the state of Amazonas, Brazil, were associated with three types of river coloration: white, black and mixed. The results demonstrated the existence of a trend towards a decrease in malaria over the years, in all types of river coloration, and that this trend does not seem to be associated with the different types of water. However, the research indicates that places located near black-or mixed-water rivers have a higher incidence of malaria, when compared to places on the banks of white-water rivers.

## Introduction

Malaria is a disease that has a great impact on morbidity in tropical and subtropical countries and, in 2020, 241 million cases were registered worldwide [1]. In Brazil, the geographic distribution of the disease is heterogeneous, with variations over time and space, and areas with a high incidence rate and regions that are free of malaria or that have a low risk of transmission can be found [2,3,4]. Most of the cases registered in the country are concentrated in the northern region, with the state of Amazonas having the highest incidence of the disease. In areas outside the Amazon region, autochthonous transmission is practically non-existent and more than 80% of cases registered are originate from the states in the endemic area or from other Amazonian countries [5,6].

Malaria is an infection caused by parasites of the genus *Plasmodium*, which are transmitted to humans via the bite of the female *Anopheles* mosquito. The main vector of malaria in Brazil is *Anopheles darlingi*. The mosquito normally reproduces on the margins of water bodies, with low flow and low depth, usually in clean waters, such as ponds, dams and ditches, in partially shaded places with superficial vegetation and with low levels of salts and organic matter [5,7,8]. However, in high-density situations, the mosquito occupies several other types of breeding sites, such as small accumulations of water [7,5].

A variety of factors can contribute to the occurrence of malaria outbreaks and epidemics, including ecological conditions, poor sanitation, climatic conditions, environmental degradation and hydrological conditions [3,8,9,10,11,12,13,14]. Rivers play an important role in the life cycle of malaria, since the vector reproduces in an aquatic environment [15]. The waters of the rivers of the Amazon have different chemical characteristics from other regions of the country, which is particularly due to the geology of the region, the type of vegetation, the presence of decomposing organisms and the climate [16,17,18]. Suspended sediments in rivers, which are generally sand, clay particles and silt, are one of the factors that determine the color of their waters. In the Amazon, river colors are classified as either white-, black-or clear-waters [16].

The white-water rivers were so named due to their origins in the Andes Mountains. They transport a large amount of suspended sediments and have a pH that is close to neutral. Among them, the Madeira, Purus, Juruá and Solimões/Amazonas Rivers stand out. Black-water rivers, on the other hand, get this name because of their characteristic color, which results from the substances dissolved in them. Humic and fulvic substances stand out above all. These rivers have an acidic pH, carry a lot of organic matter and have a low concentration of suspended sediments in their waters. The largest of these is the Rio Negro, which originates between the Orinoco and Amazon River basins. Finally, the clear-water rivers (e.g., the Tapajós River) have or greenish coloration or are transparent. These rivers carry a small amount of dissolved sediments, have a certain level of dissolved organic matter and a pH that is close to neutral.

In this context, this study correlates the colors of water in rivers in the Amazon and the incidence of malaria in order to verify whether the characteristics of the type of water can favor the life cycle of the malaria vector and, consequently, influence the distribution pattern of the disease.

## Methods

Located in the northern region of Brazil, Amazonas is the largest state in the country and has an area of 1,559,167.878 km^2^. Although, in 2021, it had an estimated population of 4,269,995 million inhabitants that are distributed in 62 municipalities, the state of Amazonas has one of the lowest indices of demographic density in the country (2.23 inhabitants/km^2^, in 2010) [19].

The studied region has an extensive network of rivers and is within the largest hydrographic basin in the world, the Amazon basin. Of the 62 municipalities of the state of Amazonas, 50 municipalities were included in this study (Fig 1) due to the availability of data for the variables studied, for being municipalities located on the banks of white-water or black-water rivers. We opted for the exclusion of municipalities near clear-water rivers due to the low number of them in the state of Amazonas.

**Fig 1:**
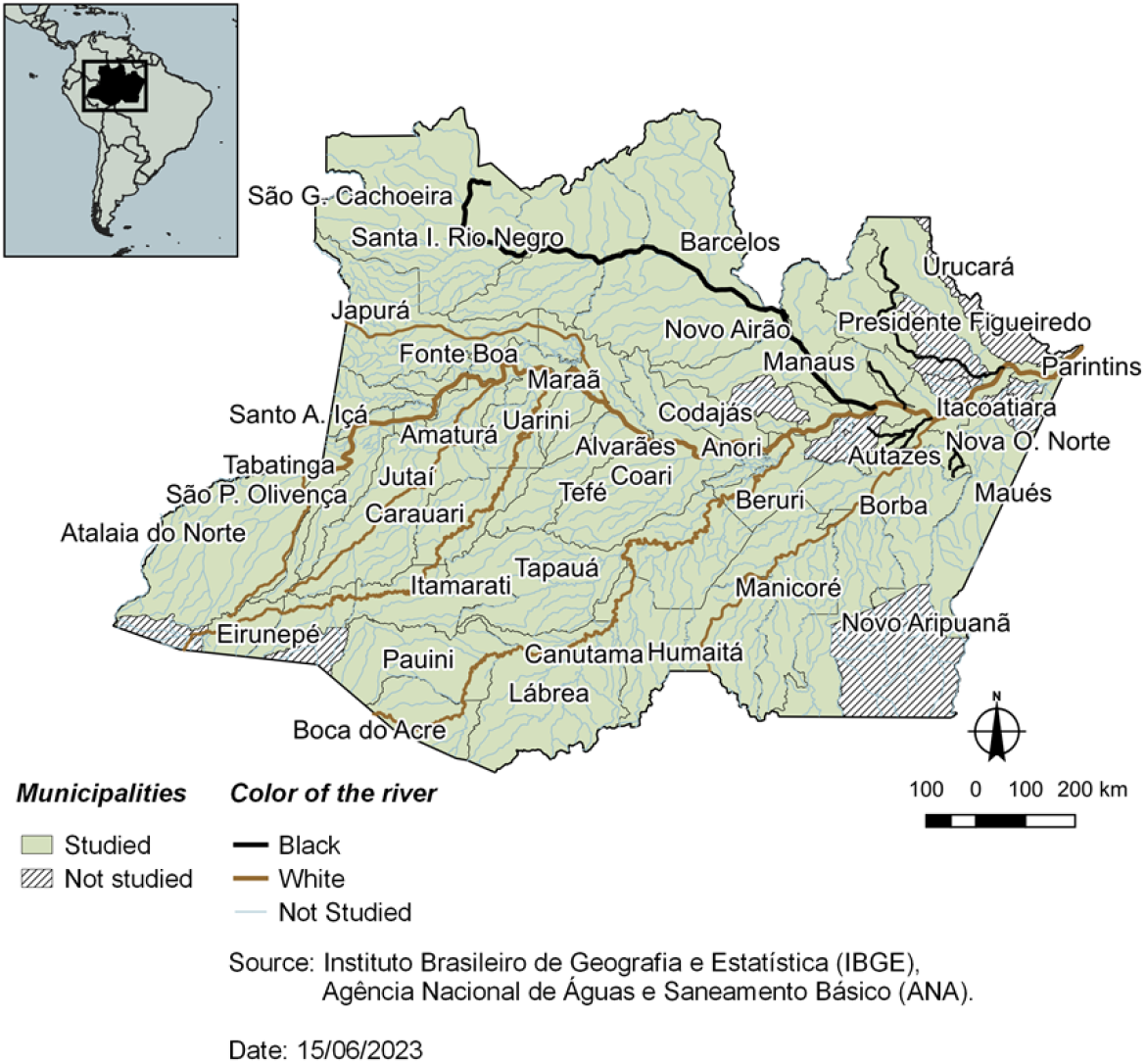
Area of study - Amazonas state, Brazil.

In addition to rivers with waters of white or black coloration, in certain stretches of the rivers, there is the mixture of both colors. These watercourses were classified here as being of mixed waters. Due to the difficulty in spatially representing the beginning and end of mixed waters, this type of water class was not depicted in Fig 1; however, they were considered in the characterization of the type of water colorations for each municipality investigated.

For the identification of the color of the rivers, in addition to the visual interpretation carried out via satellite images using the Google Earth application, information from fluviometric stations belonging to the national hydrometeorological network (www.snirh.gov.br/hidroweb) and the database of the Observatory for Environmental Research on the Hydrology and Geodynamics of the Amazon Basin, ORE/HYBAM (www.ore-hybam.org) was also used. The classification of the color of the waters of the rivers was based on the river where the fluviometric station closest to the headquarters of each municipality is found, in other words, the river closest to areas of the greatest human occupation.

Data on reported cases of malaria were obtained from the SIVEP-Malaria system, which is available to authorized users at http://www.saude.gov.br/sivep_malaria. Due to having digital records of the disease only from the year 2003 onwards, the historical series used in the study corresponds to the notifications of cases of malaria caused by *Plasmodium vivax* or *Plasmodium falciparum*, according to (most-probable) municipality of infection, between the years 2003 and 2019. The research met the criteria of resolution No. 466/2012 of the National Health Council and was approved by an ethics committee in research involving human beings.

For comparison of malaria data between municipalities, and in order to evaluate the influence of the different types of coloration of the rivers in the Amazon, the data were standardized by calculating the annual parasitic incidence of malaria (API), as detailed in [20], whose formula is as follows:

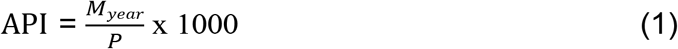

In which *M*_*year*_ corresponds to the number of positive malaria tests, excluding the cure verification slides (CVS), in the year, by probable municipality of infection, and *P* refers to the total resident population in the year considered. The CVS can be deleted, if necessary, since they represent slides of the same person. The API is a commonly used indicator to analyze annual variations in positive laboratory tests for malaria in endemic areas, as part of the group of epidemiological surveillance actions for the disease.

To assess whether the incidence of malaria is associated with the coloration of the rivers, a generalized linear mixed model (GLMM) was developed for the 50 municipalities of the study. The GLMM, proposed in [21], is made up of regression models that encompass fixed and random factors. In the analysis, fixed factors are shared by all the individuals, while random factors are specific to each. The use of the random effect in the model allows one to separate the variation between groups of individuals from the internal variation of each group of individuals, thus providing estimates of individual trajectories in time [21].

For the construction of the GLMM, the API was assigned as a response variable in the model and, as explanatory variables, the year of incidence of the disease and the color of the rivers were used, the latter being a categorical variable of classes: white-, black- and mixed-waters. The 50 municipalities investigated in the study were implemented in the model as random effects.

The GLMM (Equation 2) was generated from 850 observations, which correspond to the time series from 2003 to 2019, for the 50 municipalities, with the following structure:

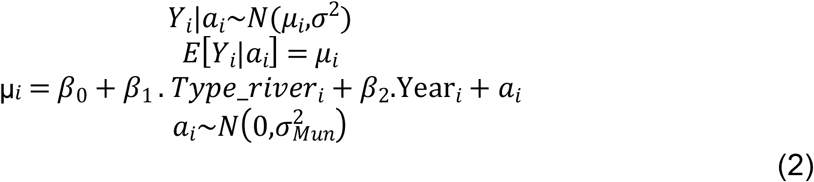

The annual parasitic incidence of malaria (*Y*_*i*_) in the municipality *i* = 1,…, 50 is explained by the fixed effect *β*_1_.*T*ype_*river* added to the year of occurrence *β*_2_.*Year*, and by the random effect a_*i*_, which is added to the value of the fixed intercept of the model, *β*_0_ to form the intercept of the municipality *i*.

The analyses were performed in the R software (version 4.1.0) in the RStudio development environment (version 1.4.17). As a decision-making tool, the significance level used in all statistical tests was 0.10. In the search for a better fit of the model, logarithmic transformation in the response variable and smoothing of the variable referring to the year of incidence of the disease were performed using the B-Spline (BS). Diagnostic analysis of residuals was performed, with analysis of residual plots versus adjusted values and envelope plots (half-normal plot).

## Results

The GLMM (Equation 2), developed in the R software and reproduced in Equation 3, comprises 850 observations, which correspond to the time series of the 50 municipalities investigated.

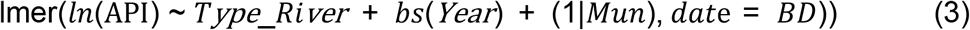

Where *ln*(API) is the logarithm of the response variable annual parasitic incidence of malaria; *Type_river* is the color of the waters of the rivers that cross the state of Amazonas, this being a categorical variable, and which are classified as white, black and mixed; *bs*(*year*) indicates the smoothing of the variable year, which expresses the year of occurrence of the case of malaria and *Mun* corresponds to the 50 municipalities that were incorporated in the regression model as a random effect.

Table 1 shows the results of the estimation of the regression coefficient (*β*), confidence interval (CI) and p-value (with accepted significance up to 0.10), generated by the GLMM.

**Table 1:** Results of the estimation of the GLMM.

| Variable | Estimate | CI | p-value |
| --- | --- | --- | --- |
| (Intercept) | 2.84 | 2.42 – 3.26 | <0.001 |
| Mixed water | 0.72 | 0.06 – 1.38 | 0.07 |
| Black water | 0.90 | 0.16 – 1.64 | 0.05 |
| Conditional R<sup>2</sup> |  | 0.66 |  |
| Reference<br>(Type_river) |  | White water |  |

In the model presented in Table 1, the category “white water” was used as the reference category of the variable color of rivers (Type_river) in relation to the categories “mixed water” and “black water”. The model produced explained 66% (conditional R^2^) of API variability and obtained a good fit to the data. Analyzing the p-value, it is observed that the categories mixed water and black water presented statistical differences in relation to white water (p = 0.07 and p = 0.05). In other words, according to the data obtained, there is evidence that the color of river waters may influence the dynamics of malaria. When observing the values of the estimate of both categories, positive values are noted (mixed water: 0.72 and black water: 0.90), which indicates that places located near mixed-or black-water rivers have a higher incidence of malaria when compared to municipalities on the banks of white-water rivers.

To better understand the analysis of the influence of river coloration on malaria incidence over time, Fig 2 presents the result of the GLMM, with the influence on API and of each category of water color type, also considering the marginal effect.

**Fig 2:**
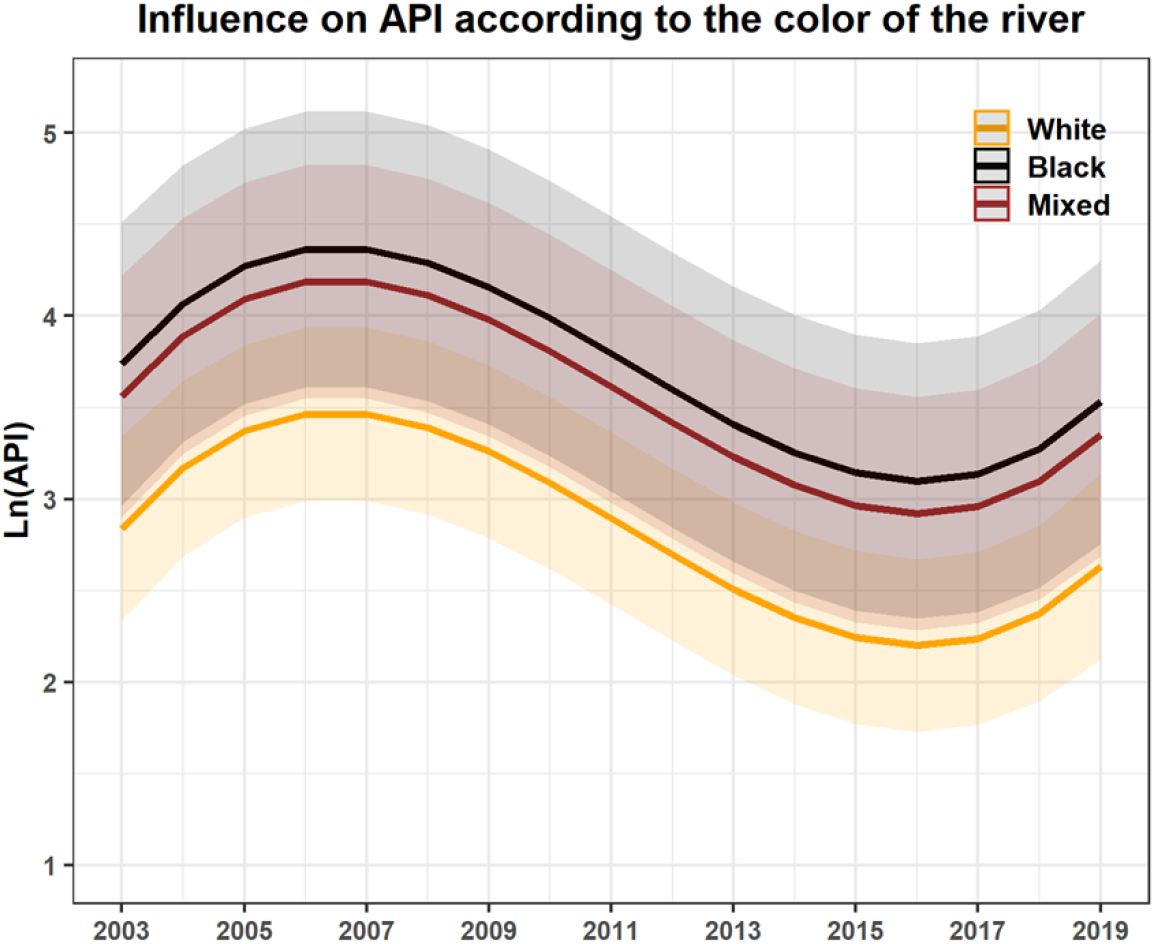
Result of the GLMM: influence on annual parasitic incidence (API) of each of the types of the Amazonian rivers (white, black and mixed water).

The results suggest that there is a trend towards a decrease in cases of malaria until the year 2015, with a possible resumption in the incidence of the disease beginning in 2017, in all types of river coloration. This trend does not seem to be associated with the different types of water, since the decrease and increase are constant in all categories.

On the other hand, the results indicate that the influence on API is differentiated between the coloration of the rivers. The results show that municipalities that are located on the banks of mixed-water rivers have more influence on the API when compared to those on the banks of white-water rivers. The incidence of malaria in mixed-water municipalities is 0.72 units (25.7%), which is higher than in that of white-water municipalities. This pattern is even clearer in the comparison between black-water rivers and white-water rivers, because our results indicate that when municipalities are located on the banks of black-water rivers there is an even more evident effect on API when compared to those of white-water rivers. The incidence of malaria in the municipalities of black-water rivers is of 0.90 units, which is equivalent to a 32% greater influence on API when compared with the API in municipalities of white-water rivers.

## Discussion

The decrease in malaria over the years has also been reported in other studies in the Amazon and around the world [1,22,23]. In recent decades, government initiatives have driven the progress in combating malaria, and this has resulted in reductions in the incidence of the disease and, thus, reductions in severe cases and deaths. Since 2000, there has been a significant increase in the number of countries that have achieved the elimination of malaria [24]. As has occurred in the global scenario, the downward trend in cases of the disease in Brazil can be explained by the prioritization in the national health policy agenda regarding the need to prevent a sharp increase in the disease. In the state of Amazonas, in 2007, there was a large investment in malaria prevention, control and surveillance actions, which boosted the progress in reducing malaria cases in the state. At that time, the multi-annual plan of malaria control actions in the state of Amazonas (PPACM 2007/2010) was implemented, which was coordinated by the Health Surveillance Foundation (HSF). According to the HSF, the PPACM established the agreement between the municipalities of the state regarding the goal of achieving, by 2010, an 80% reduction in malaria compared to 2007. Great success was achieved during this period [22].

The hydrological cycle has shown a fundamental role in relation to the dynamics of malaria [4,15]. Our results reveal that the type of coloration of river waters is a relevant factor that contributes to the incidence of the disease. Other studies have also reinforced the influence of the color of the rivers in the Amazon on the incidence of malaria [2], indicating that, in addition to the oscillation in the level of the rivers, the difference in seasonality of the disease may be linked to the color of the waters of the rivers, with a more marked seasonality in municipalities on the banks of white-water rivers. In municipalities on the banks of black-water rivers, seasonality was not very evident. The authors also point out that malaria is more intense in places and periods of the hydrological cycle with low concentrations of suspended sediments in rivers, this being one of the appropriate conditions for the development of the disease vector cycle.

The findings of this work also agree with [25] and [26]. These authors indicated that the Negro River, which has black, acidic waters, low productivity, low concentrations of suspended sediments and low electrical conductivity, provides a favorable environment for the development of *Anopheles darlingi*. Suspended sediments in rivers are one of the factors that modify the physicochemical characteristics of waters, and can cause a decrease in water temperature and an increase in turbidity, pH and conductivity. According to [27], the Negro River has low values of suspended sediment concentration throughout the year. These concentrations represent less than 0.1% of those found in the Solimões River. In addition, the Negro River presents values of other parameters that are also lower when compared to the Solimões River, such as speed (0.3 vs. 1 m/s), conductivity (8 vs. 80 μS/cm at 25°C), turbidity (5 vs. 80 NTU) and pH (5.5 vs. 7.0); and higher values for temperature (1 °C) [27]. Thus, it is suggested that the characteristics of black waters when compared to white waters may provide even more suitable conditions for the presence of the malaria vector, which influences the incidence of the disease in places on the banks of these rivers.

Although the trends indicated in the results of this study and the method and analysis indicated a good relationship with other studies discussed above, it is necessary to highlight some limitations. The use of secondary malaria notification records, which are subject to underreporting and often have poor quality of records, are some of the limitations that deserve attention. Secondly, the analysis was not adjusted for variables that may be distorting the interpretation of the results. In addition, the categorization strategy of the data referring to the variable coloration of the rivers was generalized due to the operational difficulty of classifying these waters in municipalities that have a significant geographical extension and hydrographic particularities that are typical of the largest hydrographic basin in the world.

Nonetheless, this study presents an unprecedented approach regarding the analysis of water color and malaria incidence. In addition, it was carried out in a region with hydrographic characteristics that were heterogeneous enough to allow an analysis that contrasted different colors of the rivers and covered almost the entire state, which has the highest incidence of malaria in Brazil. Thus, it is believed that these results may contribute to the more precise planning of actions aimed at disease control.

## Conclusions

The study sought to analyze whether the color of the waters of the rivers in the Amazon influence the distribution of malaria. The results showed a decreasing trend of the disease over the years from 2003 to 2019, in all types of rivers (white, black and mixed water). In addition, the research indicates that places located near black-or mixed-water rivers have a higher incidence of malaria when compared to places on the banks of white-water rivers.

## Acknowledge

The authors would like to acknowledge Fundação de Vigilância em Saúde do Amazonas (FVS) for providing access to the malaria data. We also appreciate the support of the Instituto Leônidas e Maria Deane) (ILMD - Fiocruz Amazônia), the Universidade Federal do Amazonas (UFAM), the Institut de Recherche pour le Développement (IRD/França) and the Pós-Graduação em Clima e Ambiente of the Instituto Nacional de Pesquisas da Amazônia and Universidade do Estado do Amazonas (PPGCLIAMB (INPA/UEA)).

